# A paradoxical misperception of relative motion

**DOI:** 10.1101/2024.06.04.596708

**Authors:** Josephine C. D’Angelo, Pavan Tiruveedhula, Raymond J. Weber, David W. Arathorn, Austin Roorda

## Abstract

Motion perception is considered a hyperacuity. The presence of a visual frame of reference to compute relative motion is necessary to achieve this sensitivity [Legge, Gordon E., and F. W. Campbell. “Displacement detection in human vision.” *Vision Research* 21.2 (1981): 205-213.]. However, there is a special condition where humans are unable to accurately detect relative motion: images moving in a direction consistent with retinal slip where the motion is unnaturally amplified can, under some conditions, appear stable [Arathorn, David W., et al. “How the unstable eye sees a stable and moving world.” *Journal of Vision* 13.10.22 (2013)]. In this study, we asked: Is world-fixed retinal image background content necessary for the visual system to compute the direction of eye motion to render in the percept images moving with amplified slip as stable? Or, are non-visual cues sufficient? Subjects adjusted the parameters of a stimulus moving in a random trajectory to match the perceived motion of images moving contingent to the retina. Experiments were done with and without retinal image background content. The perceived motion of stimuli moving with amplified retinal slip was suppressed in the presence of visual content; however, higher magnitudes of motion were perceived under conditions with no visual cues. Our results demonstrate that the presence of retinal image background content is essential for the visual system to compute its direction of motion. The visual content that might be thought to provide a strong frame of reference to detect amplified retinal slips, instead paradoxically drives the misperception of relative motion.

## Introduction

The human visual system is exquisitely sensitive to detecting relative motion. Judging relative motion requires a world-fixed reference object near the moving target. In a study exploring this discriminative ability, Legge and Campbell (1) found that in conditions with frames of references, displacement thresholds were as low as 0.3 arcminutes. By comparison, the center-to-center spacing of foveal cones is approximately 0.5 arcminutes (2). Detecting relative motion is therefore considered a hyperacuity because the detection thresholds transcend the sampling limits of the photoreceptors (3). In the absence of a frame of reference, motion judgements are impaired (1, 4).

Yet, there are at least two reports of a special condition under which humans fail to accurately detect relative motion. Riggs et al. (5) used an optical lever technique to either stabilize the retinal image or to amplify the retinal slip. They reported that under the amplified conditions, stimuli appeared to be “locked in place.” The authors never quantified the perceived motion under these amplified conditions nor did they ever do follow-up studies on this observation. Years later, Arathorn et al. (6) found something similar when they used a system (an earlier version of the system used in this report) to directly project images onto the retina in a retina-contingent manner. They found that under conditions where the retinal slip was unnaturally amplified, the retinal image appeared stable, despite the presence of unavoidable, world-fixed retinal background content that might have served as a frame of reference.

This so-called “illusion of relative stability” for all images moving in the direction of retinal slip, regardless of amplitude, suggests that the visual system knows its direction of motion and that anything moving in a direction consistent with the direction of retinal slip, is rendered in the percept to be stable. How is the direction of eye motion determined? Specifically, is world-fixed retinal image background content needed to compute the direction of eye motion or are non-visual cues (e.g. efferent copy) sufficient? Arathorn et al. (6) were unable to answer these questions because they could not adequately control or remove world-fixed retinal content from the visual scene. We have addressed that limitation using an updated system wherein we can fully control or remove retinal image content. We designed a method of adjustment experiment to quantify the perceived motion of stimuli moving in different directions with respect to eye motion in conditions with and without retinal image background content.

Our findings indicate that the sensory signals that inform the visual system about its direction of motion are retinal-image based. In conditions with retinal image background content, the perceived magnitude of motion of images moving with amplified retinal slip was significantly lower than that of images moving in the same direction as eye motion. When we performed the same experiments with no visual content, we found that these perceptions reversed; images with increased retinal slip were perceived to have a high magnitude of motion while images moving in the same direction were perceived as having little to no motion.

The retinal background content *−* that would normally be considered to provide a strong frame of reference for detecting motion *−* paradoxically drives the misperception of relative motion.

## Results

We measured the perceived motion of stimuli moving with increased retinal slip (Gain *−*1.5) and moving in the same direction as retinal motion (Gain +1.5). The definition of the Gains are shown in Fig. 1b. Because the motion of these stimuli is dependent on retinal motion, we call these “retina-contingent stimuli.” The specific Gains were chosen for two reasons: First, because they both generate the same amount of world motion for any given retinal motion and second, because if we chose Gain +1 and *−*1, the Gain +1 condition, being stabilized on the retina, would quickly fade from view and preclude reliable matching. We additionally tested world-fixed stimuli (Gain 0) as a control. Motion perceptions were recorded under two conditions: in the presence of rich retinal image background content (Fig. 1d) and with no visual cues (Fig. 1e). In this study, we refer to these conditions as “background-present” and “background-absent,” respectively.

**Fig. 1.**
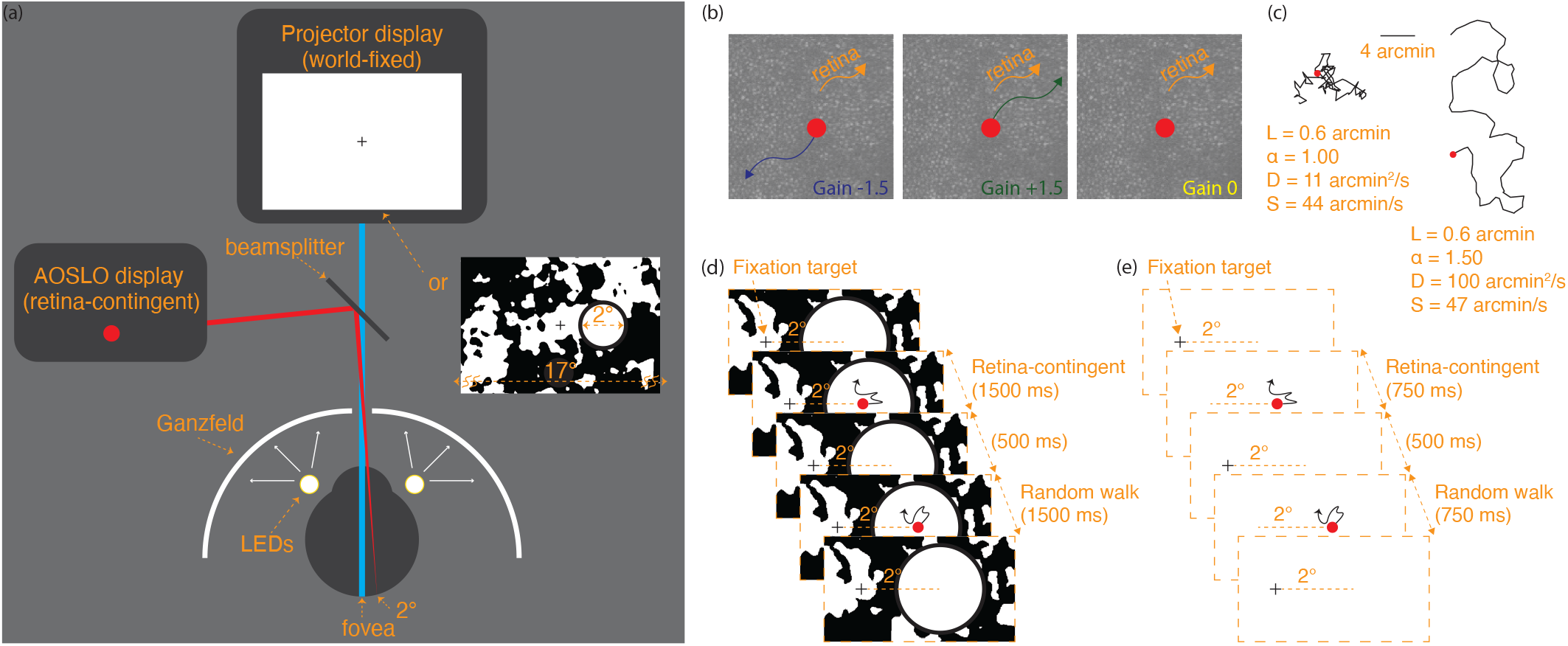
Display configuration. (a) The projector drew a fixation target and the surrounding 17° field displayed either white light or a binarized noise pattern. The AOSLO generated the stimuli positioned 2° temporally. These images were simultaneously projected onto the retina through a beamsplitter. (b) Three stimuli conditions: a Gain *−*1.5 stimulus moves with increased retinal slip, a Gain +1.5 stimulus moves in advance of retinal motion, and a Gain 0 stimulus is world-fixed. Undergoing the same retinal motion, a Gain *−*1.5 stimulus moves 2.5x more on the retina than a Gain 0 stimulus, and a Gain 0 stimulus moves 2x more on the retina than a Gain +1.5 stimulus. (c) Simulated trajectories showing uncorrelated, Brownian motion (left) and positively correlated, persistent motion (right). The red dot indicates the starting position for each trace. Both paths were generated from the same average step length, L = 0.6 arcminutes, and have similar speeds (S), yet have different diffusion constants (D) due to different *α* values. Full descriptions of these parameters are in Materials and Methods. (d-e) Experiment sequence. Subjects fixated on the target and attended to the stimuli positioned 2° temporally. The retina-contingent stimulus moved contingent to each subject’s idiosyncratic fixational eye motion. The random walk stimulus moved independent to eye motion, with pre-generated random offsets. Under background-present conditions (d), the projector field displayed a binarized noise pattern which changed after each presentation and the fixation target remained on for the entire duration. Under background-absent conditions (e), the “Ganzfeld effect” was achieved by setting up a white paper with an aperture in front of the display permitting only the AOSLO and projector beams to enter the eye, shown in (a). LEDs were taped around the eye to illuminate the paper and the luminance was adjusted so that the subject saw only the stimuli in a white full-field surround. Owing to its proximity to the eye, the natural blur of the aperture rendered the transition between the display and luminance-matched paper invisible. The fixation target was timed to turned off when the stimuli were presented.

The experiment ran as follows (full details in Materials and Methods). Subjects simultaneously viewed a world-fixed projector display and a second display, delivered through an adaptive optics scanning light ophthalmoscope or AOSLO (Fig. 1a). For each presentation, the subject held their gaze on a projector-generated fixation cross while attending to the motion of an AOSLO-generated circular stimulus presented 2° in the horizontal periphery. The retina-contingent stimulus was presented in the first time interval. Following a 500-ms insterstimulus interval, the stimulus in the second interval moved with pre-programmed random walk offsets, independent of eye motion, with a controllable magnitude quantified by its diffusion constant, D (see Materials and Methods and equation 1).

The subject’s task was to observe the motion in each interval and then adjust the D of the stimulus in the second interval (random walk stimulus) until its motion looked equivalent to that of the stimulus in the first interval (retina-contingent stimulus). Subjects could view as many self-initiated presentations as they wished for each Gain condition while adjusting the match. They were instructed to view their final match at least 3 times before submitting. The D of the random walk stimulus which the subject submitted as a match in perceived motion (PM) will be called *D*_*P M*_. All the Gain conditions were presented in pseudorandom order until six matches were submitted for each.

The AOSLO is a custom-built device that can image and track the retina in real time and deliver the retina-contingent stimuli with sub-arcminute accuracy (7). An AOSLO video of the retina containing a version of the stimulus embedded as a decrement on each video frame was recorded for each presentation. The videos were analyzed offline to compute the following: a continuous eye motion trace; the trajectory of each retina-contingent stimulus; and the accuracy of the retina-contingent stimulus. We computed the diffusion constant for the eye motion (EM) (*D*_*EM*_) and the diffusion constant for the world motion (WM) of the stimulus (*D*_*W M*_). Also, since the eye does not always exhibit true Brownian motion, we computed a second parameter, *α*, which quantified the extent to which the eye motion (and consequently the retina-contingent stimulus motion) was either persistent (*α >*1) or antipersistent (*α <*1) (8). Across all traces for each trial, we computed a single *D*_*W M*_ and *α*_*W M*_ per match, from the average of all valid retina-contingent presentations. Lastly, we computed the drift speed (*S*_*EM*_). Full descriptions of the eye motion parameters are in Materials and Methods.

Fig. 2 plots individual subject responses from experiments tested under background-present and background-absent conditions. The retina-contingent stimulus’ perceived motion (*D*_*P M*_) is plotted as a function of its world motion (*D*_*W M*_). The data points that lie along the diagonal 1:1 line represent trials where the perceived motion of the retina-contingent stimulus was equivalent to the stimulus’ actual motion occurring in the world. The red arrows depict the extent to which *α*_*W M*_ deviated from Brownian motion. Arrows pointing right represent persistence, arrows pointing left represent antipersistence, and the absence of an arrow indicates that the *α*_*W M*_ was within 1 +/*−* 0.02.

**Fig. 2.**
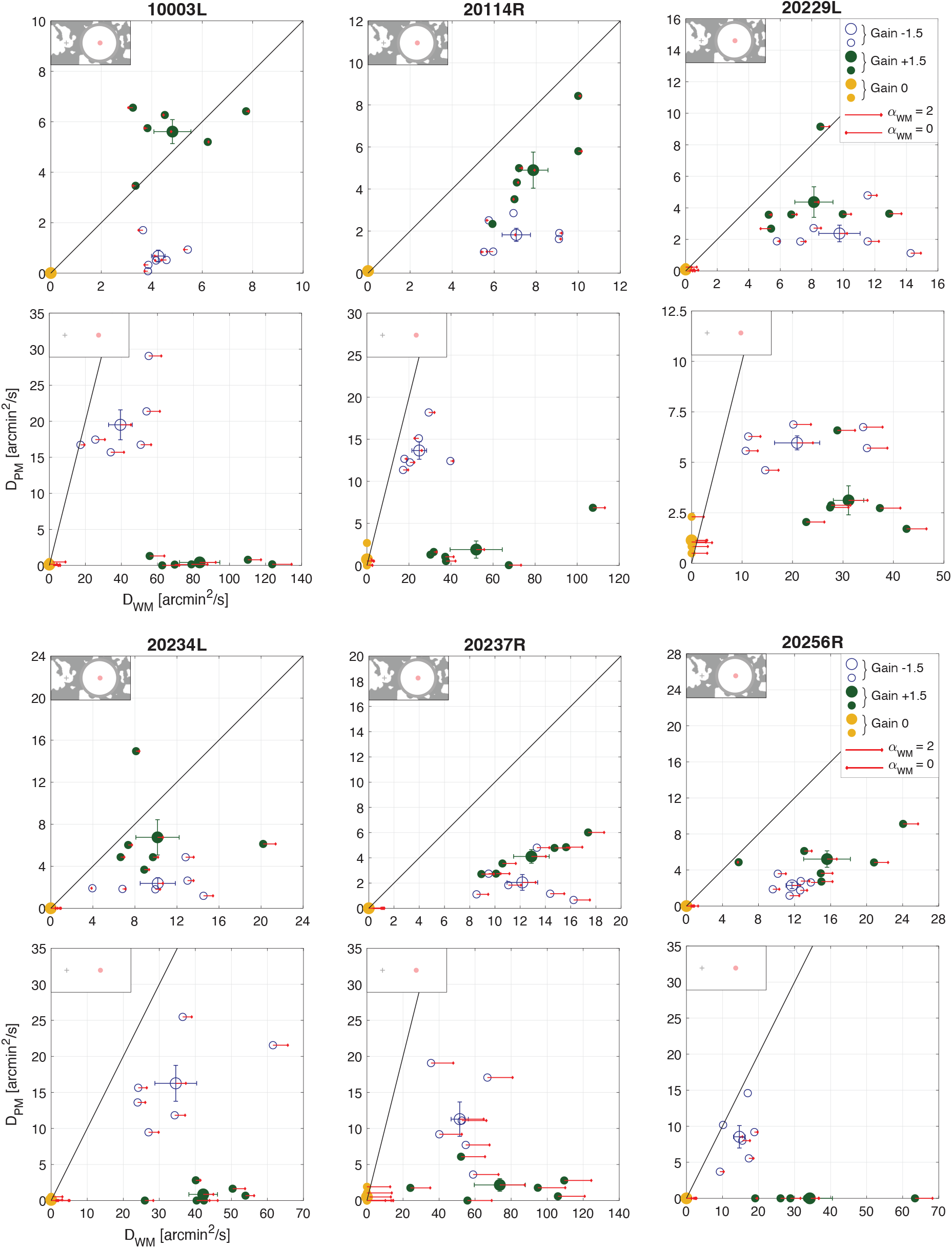
Individual results from six subjects tested under background-present and background-absent conditions. The conditions are indicated by labels on the top left corner of each graph. The small circles are the six perceptual matches for Gain *−*1.5 stimuli (blue open symbols), Gain +1.5 stimuli (green filled symbols), and Gain 0 stimuli (yellow filled symbols). The large circles are the averages of the matches with standard error of the mean bars. The red arrows show the extent to which the *α*_*W M*_ deviated from Brownian motion. Arrows pointing right indicate *α*_*W M*_ *>* 1, arrows pointing left indicate *α*_*W M*_ *<* 1, and no arrow means that *α*_*W M*_ = 1 +/*−* 0.02. Longer arrows correspond to higher deviations from Brownian motion. The arrow length in the legend indicates pure persistence (*α*_*W M*_ = 2) if pointing right or pure antipersistence (*α*_*W M*_ = 0) if pointing left.

### A. Comparing perceived motion for Gain +1.5 and *−*1.5

Under background-present conditions (Fig. 2), images moving in the same direction as eye motion (Gain +1.5) were perceived to have more motion than images moving with increased retinal slip (Gain *−*1.5). This means that despite the Gain *−*1.5 stimuli having similar world motion, as well as approximately five times more retinal motion than the Gain +1.5 stimuli, all subjects reported seeing it as having significantly less motion (p = 0.0029, post hoc Tukey-Kramer). This is consistent with the findings of Arathorn et al. (6) for which some world-fixed background content could not be avoided.

Under background-absent conditions (Fig. 2), the motion perceptions reversed. Images moving with increased retinal slip (Gain *−*1.5) were now perceived to be moving with a high magnitude of motion whereas subjects reported perceiving little to no motion when images moved in the same direction as eye motion (Gain +1.5).

The striking reversal in perceptions shows how profoundly the presence of retinal image background content impacts perceived motion. For Gain *−*1.5 stimuli, all subjects reported perceiving a higher magnitude of motion under background-absent conditions (average *D*_*P M*_ = 12.53 arcmin^2^*/*s, sem +*/−* 2.04 arcmin^2^*/*s) compared to background-present conditions (average *D*_*P M*_ = 1.93 arcmin^2^*/*s, sem +*/−* 0.27 arcmin^2^*/*s). Motion perceptions between these conditions were significantly different (p = 0.0051, post hoc Tukey-Kramer). For Gain +1.5 stimuli, all subjects reported perceiving a lower magnitude of motion under background-absent conditions (average *D*_*P M*_ = 1.41 arcmin^2^*/*s, sem +*/−* 0.48 arcmin^2^*/*s) compared to background-present conditions (average *D*_*P M*_ = 5.16 arcmin^2^*/*s, sem +*/−* 0.39 arcmin^2^*/*s). Motion perceptions between these conditions were also significantly different (p = 0.0052, post hoc Tukey-Kramer).

### B. Ratios analysis

To compare the *D*_*P M*_ and *D*_*W M*_ between experiments tested under background-present and background-absent conditions, we computed the ratio [*D*_*P M*_ */D*_*W M*_] for each retina-contingent condition. If subjects always perceived the stimulus’ actual motion in the world, then *D*_*P M*_ */D*_*W M*_ would be equal to one regardless of background condition. Fig. 3 shows the ratios in conditions with background-present plotted on the y-axis and the ratios in conditions with background-absent plotted on the x-axis. The results do not lie on or even close to the 1:1 line – instead the connected data points for each subject show a negative slope, indicative of the reversal of motion perception with background condition.

**Fig. 3.**
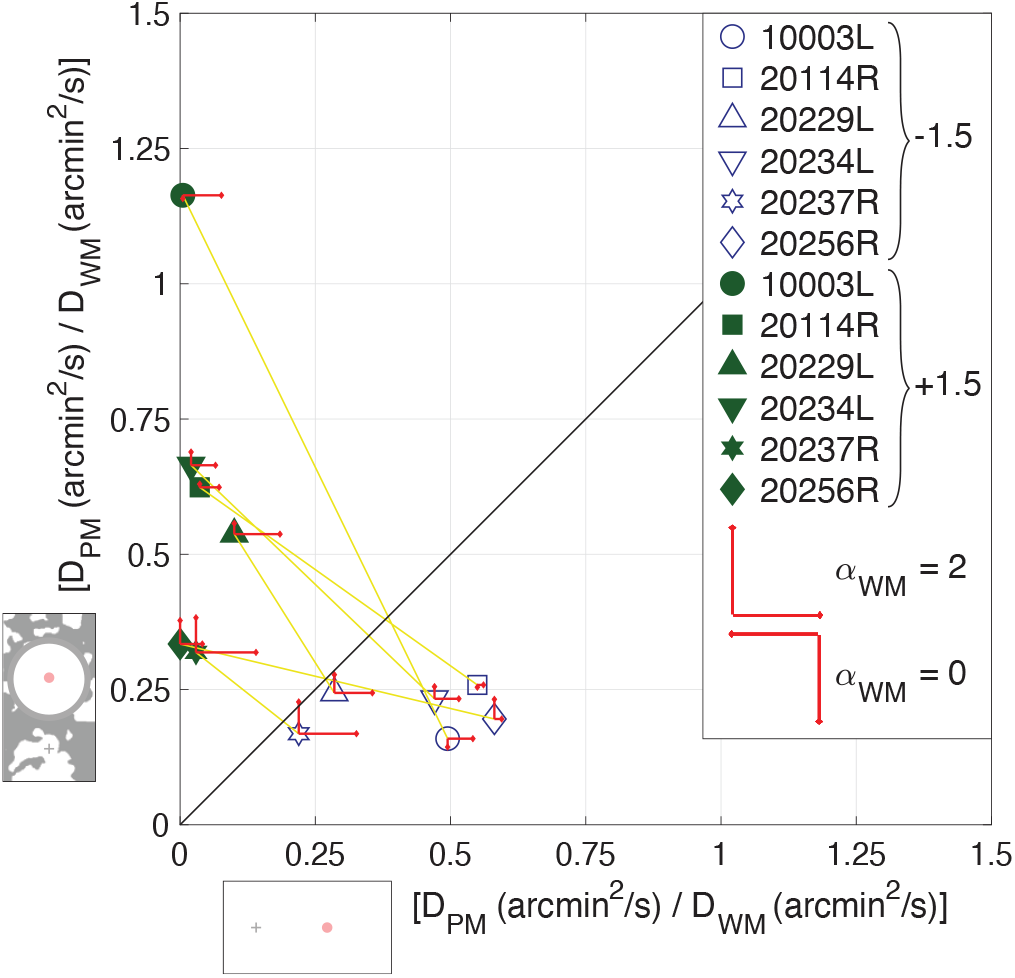
Computed ratios from six subjects tested under background-present and background-absent conditions. The red arrows show the extent to which the *α*_*W M*_ deviated from Brownian motion. Arrows pointing up and right indicate *α*_*W M*_ *>* 1 under background-present and background-absent conditions, respectively. Arrows pointing down and left indicate *α*_*W M*_ *<* 1 under background-present and background-absent conditions, respectively. No arrow means that *α*_*W M*_ = 1 +/*−* 0.02. Longer arrows correspond to higher deviations from Brownian motion. The arrow length in the legend indicates pure persistence (*α*_*W M*_ = 2) if pointing up/right or pure antipersistence if pointing down/left (*α*_*W M*_ = 0).

### C. Perceived motion for Gain 0

Under both the background-present and background-absent conditions, all subjects reported perceiving little to no motion for world-fixed images (Gain 0) (Fig. 2). Under background-present conditions the average *D*_*P M*_ = 0.0325 arcmin^2^*/*s, sem +*/−* 0.020 and under background-absent conditions, the average *D*_*P M*_ = 0.43 arcmin^2^*/*s, sem +*/−* 0.18. Motion perceptions between these conditions were not significantly different (p = 0.0642, post hoc Tukey-Kramer). For each subject, the average perceived motion for the world-fixed images was always lower than the average perceived motion of Gain *−*1.5 and Gain +1.5 stimuli.

### D. Comparing perceived motion for Gain +1.5 and 0 under background-absent conditions

Subjects reported seeing little to no motion when viewing the Gains 0 and +1.5 stimuli under background-absent conditions. Motion perceptions were not significantly different between the Gains 0 and +1.5 stimuli (p = 0.0574, post hoc Tukey-Kramer). The eye motion was not significantly different between Gain +1.5 and Gain 0 presentations, shown in Fig. 4a. However, the Gain 0 stimuli moved across the retina with greater retinal motion (RM) than Gain +1.5 stimuli: The average *D*_*RM*_ of a Gain 0 stimulus was 15.55 arcmin^2^*/*s, while the average *D*_*RM*_ of a Gain +1.5 stimulus was 5.86 arcmin^2^*/*s (see Materials and Methods).

**Fig. 4.**
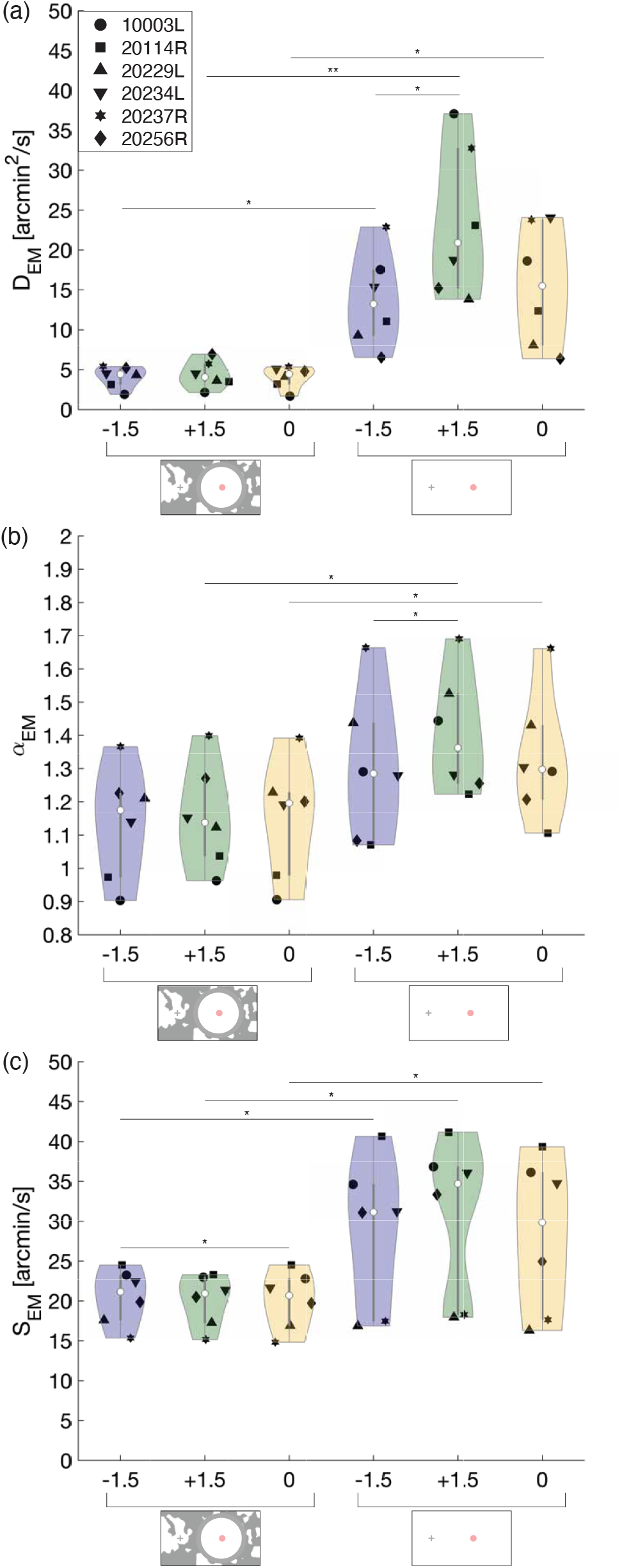
**a-c** Violin plots (9) showing the distribution of the (a) *D*_*EM*_, the (b) *α*_*EM*_, and the (c) *S*_*EM*_ for three Gains (*−*1.5, +1.5, and 0) under two conditions (background-present and background-absent). In the center of the violin, the white circle represents the median and the dark bars represent the interquartile range. The black points are the (a) mean *D*_*EM*_, (b) mean *α*_*EM*_, and (c) mean *S*_*EM*_ for each subject across their six respective trials for each Gain. The asterisks indicate statistical significance (p *<* 0.05 and p *<* 0.01, respectively) from a post hoc Tukey-Kramer test following a two-factor repeated-measures ANOVA.

### E. Eye motion depends on background and stimulus condition

The drift diffusion constants, *α* values, and speeds for all Gains and background conditions are plotted on Fig. 4. Under background-present conditions, *D*_*EM*_ was not significantly different between Gain conditions (Fig. 4a). Most, but not all, eyes exhibited a small degree of persistence (*α*_*EM*_ *>*1) but the *α*_*EM*_ did not significantly differ between Gain conditions (Fig. 4b). The *S*_*EM*_ during Gain *−*1.5 stimuli presentation was slightly higher than that of the Gain 0 stimuli presentation (Fig. 4c).

The *D*_*EM*_ and *S*_*EM*_ were uniformly greater for all Gains under background-absent conditions, meaning there was more eye motion. While *α*_*EM*_ values were greater for all Gains under background-absent conditions, only Gain +1.5 and 0 conditions showed a significant increase.

Under background-absent conditions, the *D*_*EM*_ of the Gain +1.5 stimuli was significantly greater than that of the Gain *−*1.5. The *α*_*EM*_ values were greater than one for all Gain conditions but were greatest for the Gain +1.5 condition.

To offer a physical sense of the differences in *α*_*EM*_ values between background conditions, all the eye trajectory traces during the Gain *−*1.5 retina-contingent stimulus presentations are plotted for one of the subjects in Fig. 5 a and b.

**Fig. 5.**
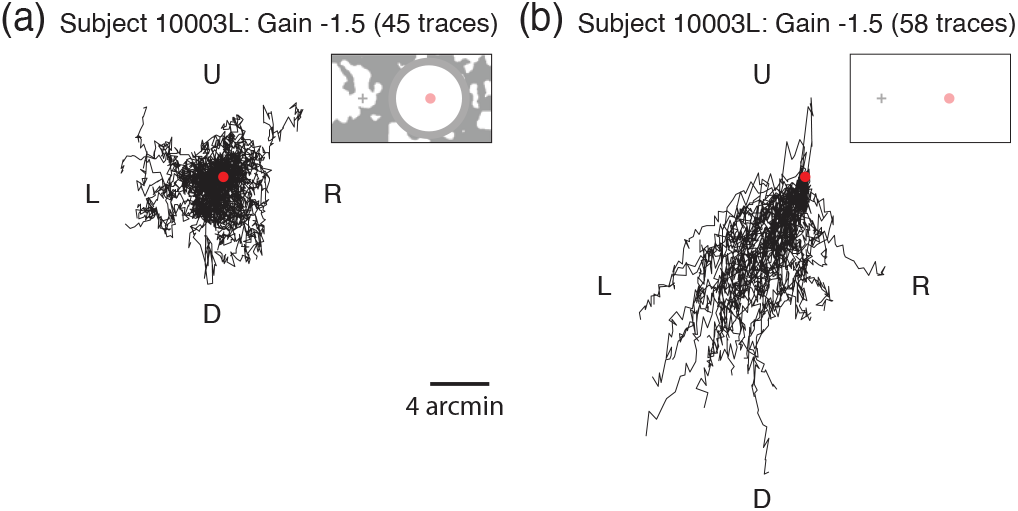
Example gaze traces for subject 10003L during Gain *−*1.5 retina-contingent presentations under (c) background-present (1500-ms duration and *α*_*EM*_ = 0.90) and (d) background-absent (750-ms duration and *α*_*EM*_ = 1.29) conditions. The red dot indicates the starting position for each trace. The gaze directions are labeled: left (L), right (R), up (U), and down (D).

### F. The presence of persistent eye motion (*α*_*EM*_ *>*1) affects interpretation of the results

Subjects adjusted the diffusion constant of the random walk stimulus until its motion looked perceptually equivalent to the motion of the respective retina-contingent stimulus. Although Fig. 4 shows that *α*_*EM*_ changed between conditions and was mostly persistent, we chose to constrain the motion of the random walk matching stimulus to be Brownian motion with *α*, on average, equal to one.

We constrained the motion of the random walk matching stimulus to be Brownian motion because we found that subjects perform poorly at discriminating a motion with a high *α* and high D from a motion with a low *α* and low D. This was discovered when we ran a control experiment with the same method-of-adjustment procedure, but with controlled random walk stimuli in both intervals. The stimulus in the first interval was a pre-programmed random walk stimulus with a selected *α* and D value. The task was to adjust the random walk stimulus in the second interval until its motion looked equivalent to that of the random walk stimulus in the first interval. Subjects could adjust two parameters – the D as well as the *α*. We repeated each match three times. We found that for the same test D and *α* values, sometimes the subject would correctly match the D and *α*, while for other matches the D and *α* values were both much higher or both much lower. What could contribute to this mismatch, is that random walks with high D and *α* values can be generated from the same average step length as random walks with low D and *α* values, examples are shown in Fig. 1c. We concluded that introducing a second adjustable parameter *α* complicated the interpretation of results and also made the experimental procedure more difficult for the subject. We therefore ran experiments with one adjustable parameter, the D, and held the *α* constant at one.

We could not, however, control the *α* of the retina contingent stimuli because the subjects’ fixational eye motion governed this parameter. When the *α*_*W M*_ is different from the *α*_*P M*_, comparing only their D values is too simplistic. This is because the D and *α* values are dependent on each other. A subject with persistent eye motion, would have a high *α*_*W M*_ value and thus a higher D value, compared to a trajectory with the same average step length but with an *α* equal to one. This could explain why, in Figure 3, the subjects with high *α*_*W M*_ did not report *D*_*P M*_ equal to the *D*_*W M*_. The subjects with more persistent *α*_*W M*_ (larger red arrows) have lower ratio values. These higher *α*_*W M*_ values correspond to higher *D*_*W M*_ which increase the value in the denominator.

Figure 6 plots the ratios from Figure 3 as a function of each subject’s *α*_*W M*_, for two of the conditions: Gain *−*1.5 stimuli tested under background-absent conditions and Gain +1.5 stimuli tested under background-present conditions. The solid line is a plot of *D*_*P M*_ /*D*_*W M*_ versus *α*_*W M*_, where the random walk stimulus’s *α*_*P M*_ is equal to one (Brownian motion) while the retina-contingent stimulus’s *α*_*W M*_ ranges from 0.9 to 1.8. As the retina-contingent stimulus’ *α*_*W M*_ becomes more persistent, it on average traverses further than a Brownian stimulus and therefore the MSD of the trajectory increases faster over a time interval compared to the Brownian stimulus. The model shows that there is an exponential decay relationship between *D*_*P M*_ /*D*_*W M*_ and *α*_*W M*_ – as the eye motion becomes more persistent (*α*_*W M*_ *>* 1), the corresponding *D*_*W M*_ increases and subsequently the ratio *D*_*P M*_ /*D*_*W M*_ decreases. The human subject data are largely predicted by this model.

**Fig. 6.**
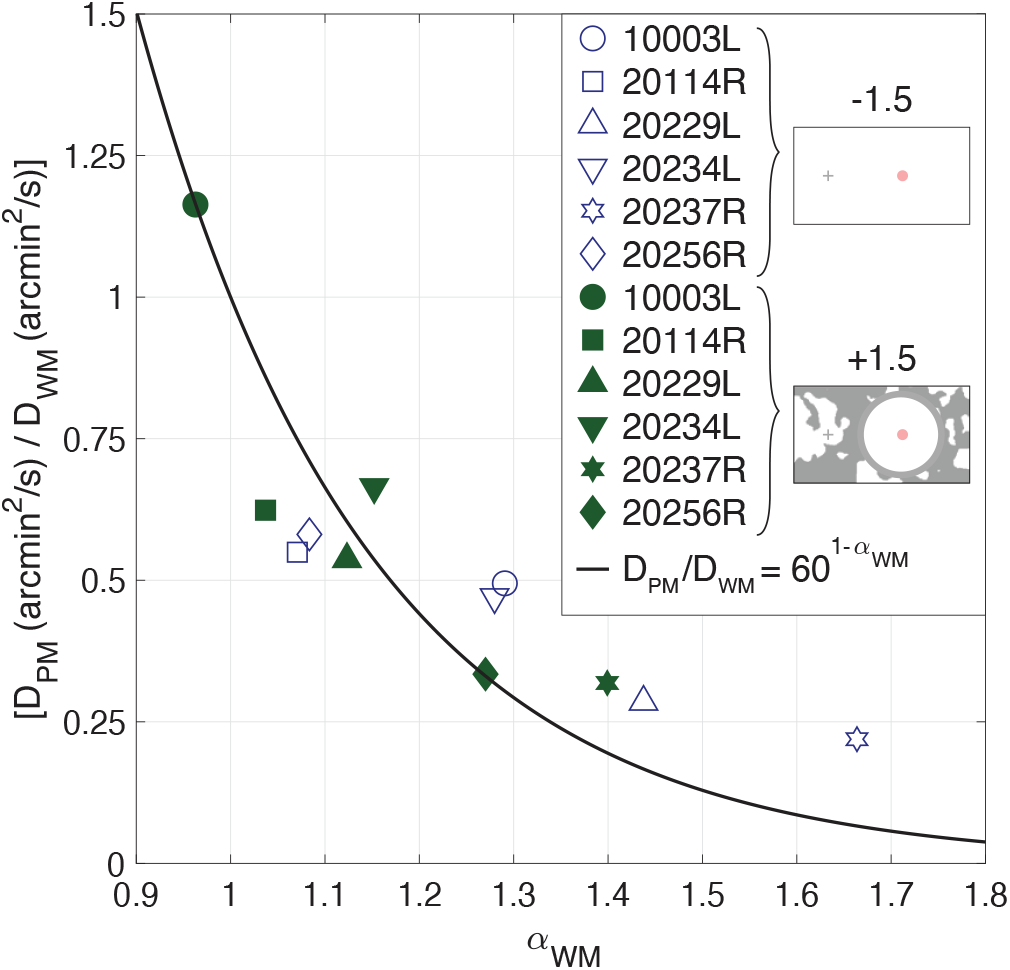
Computed ratios from six subjects plotted as a function of each subject’s *α*_*W M*_. Data from two experiments are shown: Gain *−*1.5 stimuli tested under background-absent conditions and Gain +1.5 stimuli tested under backgroundpresent conditions. The black line is the simulated *D*_*P M*_ /*D*_*W M*_ versus *α*_*W M*_, where the *MSD*_*P M*_ = *MSD*_*W M*_ but the random walk stimulus’s *α* is equal to one (Brownian motion) while the retina-contingent stimulus’s *α*_*W M*_ ranged from 0.9 to 1.8. Both the model and data points show exponential decay.

## Discussion

When an object moves in the world, the image that is cast onto the retina has retinal motion both due to the object’s motion as well as the fixational eye motion. To properly perceive an object’s motion in the world, the visual system is tasked with disentangling the object’s motion from the fixational eye motion (10, 11). Normally, the visual system performs this task exceptionally well; humans are able to reliably perceive world-fixed objects as stable and can identify moving objects within it with hyperacuity (1, 3). Yet, in conditions where an image is programmed to move with amplified retinal slip, humans exhibit a paradoxical inability to accurately perceive the motion of the image relative to a high-contrast, world-fixed background.

Our findings support those of Arathorn et al. (6) and demonstrate that the visual system suppresses the perceived motion of anything moving in a direction consistent with retinal slip (Gains less than 1), despite their magnitude. The neural mechanism for doing this has yet to be described but it is functionally sufficient, since the likelihood that any real-world object would be moving in the same direction of retinal slip for any appreciable duration is effectively zero. All other directions, such as our Gain = +1.5 and all other Gain conditions described in Arathorn et al. (6), are perceived as moving. As a control, we tested the same experiment for one subject, presenting a stimulus that moved with a Gain of +1.5, but programmed to move orthogonal to eye motion. There was no significant difference in motion perception between this stimulus and the stimulus that moved in advance of eye motion, Gain +1.5.

This paper shows that the direction of eye motion must arise from the retinal image content itself (inflow) and is not relayed to the visual system by any other means, such as efference copy.

To further understand the process, we first review what is known functionally and structurally about perception of motion.

### G. Perceiving the world as stable

Previous studies showed that the retinal input informs the visual system about image motion on the retina due to drift. Poletti et al. (12) presented gaze-contingent stimuli using a dual Purkinje eye tracker in conditions with and without frames of references. Consistent with our findings on Fig. 2, stimuli moving in the same direction as gaze (theirs was a Gain +1 condition, which meant the image was stabilized) were perceived to move when frames of references were present, and perceived as stable when all visual content was removed. They concluded that the retinal input drives the compensation of retinal motion. Based on a study of the jitter-aftereffect illusion, Murakami and Cavanagh (13) also concluded that retinal signals inform the visual system on image motion due to retinal motion.

A possible origin of the retinal signal is directionally sensitive retinal ganglion cells (DSRGCs) for which evidence of their existence in primate and human retina continues to mount (14–16). The subclass of ON-OFF DSRGCs, which project to the LGN and subsequently the visual areas of the cortex, are the most likely candidates. Combined signals from a sufficient number of these cells to eliminate ambiguity could be used to compute the direction of retinal slip, which may provide partial information to the downstream circuitry to assist in the stabilization process. Exactly how and where the stabilization occurs however, is not known and evidence is mixed (17–20).

### H. Detecting moving objects

Humans have a hyperacute ability to detect (21) and resolve (1) the motion of objects within a visual scene. The neural underpinning for detection of these stimuli in the presence of incessant eye movements may lie in the Object Motion Sensing (OMS) ganglion cells. Yet to be found in primates, this class of retinal ganglion cells is very effective at identifying an object that is moving differently than the surround (11).

But despite increasing knowledge of the neural systems that underlie our ability to perceive a stable and moving world (22), what remains unclear is how two objects that move in a direction consistent with retinal slip but with different velocities can be rendered in the percept to be fixed relative to each other.

### I. The role of efference copy

Studies have shown that efference copy or other non-visual cues can inform the visual system about its gaze direction during drift (23, 24), albeit not with great accuracy. In our experiments, however, it is very clear from the background-absent condition that neither efference copy nor any other nonvisual signal is being used effectively to determine the direction of motion in a manner that guides perception of moving objects. That being said, the perception of motion in the background-absent condition cannot be explained as being due simply to retinal motion either. For example, the Gain 0 condition, which actually moved more across the retina than the Gain +1.5 condition, was perceived to be moving about the same, or even less than the Gain +1.5 condition. Further investigations of this condition are warranted.

### J. Control experiments to validate results

In this study, we presented stimuli for 1500-ms under background-present conditions and for 750-ms under background-absent conditions. These duration choices were motivated by a trade-off to achieve robust motion judgements while also optimizing retinal tracking. A longer interval duration gives the subject sufficient time to discern the motion magnitude of the stimuli but can also lead to tracking failures if the eye drifts too far away from fixation. We performed control experiments on subjects 10003L and 20114R to verify that different stimulus durations did not contribute to the results. We presented the stimuli for 750-ms intervals under background-present conditions and found similar trends, although less robust, as experiments with 1500-ms intervals. Under background-absent conditions, the eye had faster drift and larger corrective microsaccades which led to tracking failures. As a result, we were unable to perform the experiments with 1500-ms stimulus intervals under background-absent conditions.

In the experiments tested under background-absent conditions, we ensured no frames of reference were visible to the subject, even removing the fixation target during the retina-contingent interval and the random walk interval. Under these conditions, we could not truly assess the amount of motion perceived. Specifically, we could not conclude that the subject did not perceive motion when they matched a Gain 0 condition to a non-moving stimulus in the second interval. As a check, we performed a control experiment for one subject with the same experimental protocol, except that the fixation target remained on to serve as a frame of reference during the random walk interval. We found no difference in matches to the Gain 0 stimuli whether the fixation target was on or off during the random walk interval.

This study shows the power of combining high resolution retina-tracking with stimulus delivery to reveal properties of motion perception that would otherwise not have been discovered in natural conditions. Our results provide valuable data that will shed light on and inform about the neural underpinnings of this important property of human vision.

## Methods

### K. Subjects

Six subjects (4 experienced and 2 naive) were recruited. The experiment was approved by the Institutional Review Board at the University of California, Berkeley. Prior to the experiment, subjects provided informed consent to participate. We applied topical eye drops of 1.0% tropicamide and 2.5% phenylephrine hydrochloride to dilate and cycloplege subjects prior to each experiment.

### L. Adaptive optics scanning light ophthalmoscopy

Experiments were performed using a multi-wavelength adaptive optics scanning light ophthalmoscope (AOSLO) (25, 26). A 940 nanometer (nm) laser beam measured the eye’s wave-front and a deformable mirror corrected for the optical imperfections of the eye to confine the laser beams to a small spot. A focused 840-nm laser beam scanned sinusoidally across the retina by a 16 kHz horizontal scanner and a 60 Hz vertical scanner. The field size was set to image a 1.71° raster square of the retina. An acousto-optic modulator (AOM) modulated a 680-nm laser beam to deliver a circular 12 arcminute diameter increment stimulus onto targeted retinal locations. Concurrently, the AOM modulated the 840-nm imaging beam to turn off at the same targeted retinal locations. This rendered a 3 x 3 arcminute decrement in the image which enabled unambiguous tracking of the 680-nm increment stimuli. We used custom software to move the stimuli contingent to the retinal motion (7). The AOSLO records real-time, high resolution 512 by 256 pixel videos of the retina. Each pixel subtends 0.2 by 0.4 minutes of visual angle. The average power of the 940-nm, 840-nm, and 680-nm laser beams were 58.6 *μ*W, 111.4 *μ*W and 11.6 *μ*W giving rise to equivalent luminances of 0.0036 cd/m^2^, 0.84 cd/m^2^ and 3960 cd/m^2^, respectively using methods described by Domdei et al. (27). Pupil sizes for subjects ranged from 5.9 to 7.2 mm.

### M. Projector display

A DLP LightCrafter projector (Wintech, Carlsbad, CA, USA) displayed a 17° background which projected over the 840-nm imaging raster. The mean luminance of the display, which was approximately 540 cd/m^2^, effectively canceled perception of the 840- and 940-nm rasters. A fixation target and patterns were drawn over the background and timed with the stimulus delivery dependent on the experiment conditions (Fig. 1 d and e). The procedure was programmed with MATLAB (MathWorks, Natick, MA, USA) using the Psychophysics Toolbox (28–30).

### N. Real-time eye tracking

Fixational eye movements cause distortions in the raw AOSLO videos and these distortions encode the eye motion that occurred during image acquisition (31). We extracted the eye motion traces from these videos by sectioning each frame into 32 strips, selecting a reference frame, and then cross-correlating the strips of every subsequent frame to the reference frame (7). This realtime eye tracking enabled targeted stimulus delivery; in each frame at a critical strip, the AOM would modulate the 680-nm and 840-nm laser beams to place increment and decrement stimuli at the predicted retinal location. The critical strip occurred 2 milliseconds before stimulus delivery and this reduced lag enabled subarcminute accuracy (32).

### O. Retina-contingent conditions

The high-resolution, real-time eye tracking of the AOSLO allowed us to deliver stimuli that moved contingent to fixational eye motion transformed by a “Gain.” The sign of the Gain indicates the direction the image moved with respect to the eye, and the stimulus’s world displacement was equal to the Gain magnitude times the eye displacement. We tested three Gains: *−*1.5, +1.5, and 0. The Gain *−*1.5 stimulus moved directly opposite the direction of eye motion with a magnitude that was 1.5 times that of the eye motion. For example, if the eye moved 2’ right, then the stimulus moved in the world 3’ left, producing a total retinal displacement of 5’. Therefore, in this condition, the stimulus moved on the retina with 2.5 times more motion than a natural, world-fixed object. The Gain +1.5 stimulus moved in the same direction as the eye with a magnitude equal to 1.5 times that of the eye motion. For example, if the eye moved 2’ right, then the stimulus moved in the world 3’ right. The stimulus moved in advance of eye motion with retinal image motion that was half the retinal motion of a natural, world-fixed object. The Gain 0 stimulus did not move in the world, which means that this was a natural, world-fixed stimulus and served as control.

### P. Random walks

Prior to the experiment, we preprogrammed 100s of 750-ms and 1500-ms duration random walks with varying diffusion constants (D) for the subjects to use in the matching task. These paths were generated by computing randomized step lengths in x and y drawn from a normal distribution with a range of standard deviations from 0 to 1.6 arcminutes per step (or AOSLO frame), in increments of 0.05 arcminutes. For the sake of brevity, we will refer to the standard deviation of the randomized step lengths as simply *step lengths* throughout the manuscript. This process effectively randomized the length and direction of each step and, because all directions were equally probable, the *α* of the paths was approximately equal to one. However, owing to the random nature of generating paths in this method, the same randomized step length could generate a path with a range of possible D values. We computed the average D value for each step length, by inputting the step length as the *△*X and *△*Y in the MSD formula in equation 2 which was then inputted into equation 1 to solve for the D. For example, a step length of 1.6 arcminutes corresponded to a D = 76.8 arcmin^2^*/*s. Therefore, for each of the 33 step lengths, we generated 100 paths and from these, we selected the 10 paths with Ds closest to the respective average D.

### Q. Experimental design

#### Q.1. Set up

The subject’s pupil was aligned with the AOSLO beam and their head was immobilized by the use of a dental bite bar.

#### Q.2. Conditions

We tested two conditions. Under the background-present condition, the 17° projector background was filled with rich retinal image background content: a blurred and binarized 1*/*f noise pattern that changed after every presentation (Fig. 1d). The fixation target was present for the entire duration and each stimulus was presented for 1500-ms. Under the background-absent condition, all visual cues were removed (Fig. 1e). This was achieved by attaching a white paper with a small central aperture directly in front of the subject. The aperture permitted only the raster and projector light to pass into the subject’s eye. The proximity of the aperture to the subject’s eye (about 5 cm) ensured that its edges appeared out-of-focus and blurred and therefore blended with the projector view. We then taped LEDs around the subject’s eye and the subject adjusted the power of the LEDs until the white projector light was indistinguishable from the paper. Additionally, the fixation target disappeared during the intervals with stimulus presentation. Without a fixation target, eyes generally exhibit faster drift and larger corrective microsaccades (33). To mitigate the effects of this increased retinal motion on the tracking accuracy, each stimulus was presented for 750-ms.

#### Q.3. Task

The projector display drew a fixation target positioned 2° nasally away from the AOSLO raster, which remained stationary Fig. 1 d and e. The 680-nm light from the AOSLO delivered increment circular images onto the retina, which subtended 12 minutes of visual angle. We tested one retina-contingent condition per trial, either Gain *−*1.5, +1.5, or 0. In each trial, the subject could initiate as many presentations as necessary. In a single presentation, the subject attended to the stimulus in the first time interval (retina-contingent), followed by a 500-ms break, followed by the stimulus in the second time interval (random walk). The subject’s task was to adjust the D of the random walk stimulus until its motion looked perceptually equivalent to the respective retina-contingent stimulus. When the subject found a match, they initiated a minimum of three final presentations and then submitted their response. In one block, there were six trials and the trial order was randomized. Subjects completed three blocks per background condition. Therefore, for each background condition, there were six trials tested for each Gain. All perceptual responses were made on a gamepad.

### R. Quality Control

During the experiment, if the subject looked away from the fixation target, the tracking would fail and the stimulus would not be delivered. This ensured that the subject maintained fixation when making motion judgements. Blinks and large saccades also led to tracking failures and the stimulus misdelivery was usually recognizable to the subjects, who were informed ahead of the experiment to disregard presentations with poor stimulus delivery.

After the experiments, we used custom software (34) to extract a continuous eye motion trace from each recorded video. We then used a video-analysis script to determine the retinal positions of the 3 x 3 arcminute stimulus-tracking decrement in each frame. We compared the eye motion trace to the stimulus motion trace to verify that the stimulus moved appropriately contingent to the retina, dependent on the Gain. In this way, videos were evaluated for tracking errors and stimulus misdelivery.

We determined the number of videos where the stimulus delivery was poor but might not have been recognizable to the subject. This was achieved by comparing the eye position to the stimulus-tracking decrement position in frames with retina-contingent presentation. We computed the standard deviation of the stimulus delivery accuracy for each video across all the frames that had retina-contingent stimulus presentation. We considered a video to have poor stimulus delivery when the standard deviation of the stimulus misdelivery from the targeted location was greater than 0.9 arcminutes. If the majority of the videos used by the subject had standard deviations greater than this threshold, we removed the trial. This strict criterion ensured that the motion judgements were not contaminated by presentations with poor tracking. This was especially important for the Gain *−*1.5 stimuli because tracking errors could cause stimuli motions that no longer slipped in directions consistent with retinal motion, thus disrupting the “illusion of relative stability”.

Because we are interested in motion perception during periods of fixational drift, we used a final filter that removed traces with microsaccades. Microsaccades are linear and ballistic and this would have biased the random walk analysis.

### S. Analysis of eye motion and retina-contingent stimulus motion

#### S.1. Random walk analysis

Fixational drift, the eye motion that occurs in between microsaccades, resembles a random trajectory similar to Brownian motion (8, 35–39). That is, the motion is two-dimensional with a velocity that varies according to a normal distribution. Studies have shown that drift deviates from uncorrelated random motion, instead having correlated or anticorrelated properties depending on the timescale (8, 36). To quantify the statistical properties of drift, we computed the diffusion constant (D) which quantifies the amount the eye moves from its starting position, and we computed the alpha (*α*) which measures the extent to which the steps in the trajectory are uncorrelated, positively correlated, or negatively correlated. Equation 1 shows a method to solve for D and the scaling exponent *α* (40, 41).

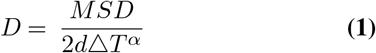

From equation 1, the D measures the temporal changes of the mean square displacement. The mean square displacement (MSD) is the squared Euclidean distance of the horizontal and vertical positions of the eye between two time points specified by the time interval (*△*T) which, in our case, is the time between frames (Equation 2). The dimension (d) is equal to two since the eye moves in both x and y directions. As suggested by Saxton (42), the number of *△*Ts was determined by one-quarter times the total time steps (in our case, total number of frames). We did not calculate beyond one-quarter of the time steps because higher *△*Ts have larger standard deviations and thus are more vulnerable to noise (42).

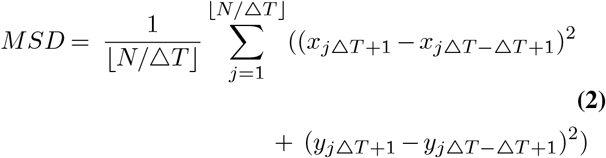

From equation 1, if *α* = 1, then the MSD increases linearly with *△*T. This is uncorrelated random motion because the steps in the trajectory are independent from previous steps. This is referred to as Brownian motion. When *α >* 1, the MSD increases faster than linearly with *△*T. The steps in the trajectory are positively correlated; each step has a tendency to continue moving in the same direction as the previous step. This is referred to as superdiffusion (43), persistence (8, 40, 44), or diffusion with flow (41). When *α <* 1, the MSD increases slower than linearly with *△*T. The steps in the trajectory are negatively correlated; each step has a tendency to move in the opposite direction as the previous step. This is referred to as subdiffusion (43), antipersistence (8, 40, 44), or caged motion (41). In this study, we use the terms Brownian motion, persistence, and antipersistence.

We can compute D and *α* by plotting the *log*_10_(MSD) as a function of the *log*_10_(*△*T). The slope of the line across these data points is *α* and the y-intercept is *log*_10_(D). Therefore, across all of the traces from each analysis, we quantified the amount of motion (D) and its deviation from Brownian motion (*α*).

We performed this random walk analysis on: the eye’s motion (*D*_*EM*_ and *α*_*EM*_), the retina-contingent stimuli’s motion (*D*_*W M*_ and *α*_*W M*_), and the random walk stimuli’s motion at the setting for a perceptual match (*D*_*P M*_ and *α*_*P M*_).

#### S.2. Speed analysis

Across all traces with retina-contingent stimulus presentation, we computed the average displacement of the eye over time to compute the speed (*S*_*EM*_). Eye positions were sampled at 60 Hz.

#### S.3. Statistics

We performed a two-factor repeated-measures analysis of variance (ANOVA) of the *α*_*EM*_, *D*_*EM*_, *D*_*P M*_, and *S*_*EM*_. There were two background conditions (background-present and background-absent) and three retina-contingent stimuli (Gain *−*1.5, Gain +1.5, and Gain 0), which were within-subject factors. Following the repeated-measures ANOVA, we performed a post hoc Tukey-Kramer test to measure the significance of the pair-wise comparisons. We compared between Gain categories within the same background condition and for each Gain we compared differences between background conditions.

### T. Comparing the eye’s motion and the retina-contingent stimuli’s world motion

The *D*_*W M*_ and *α*_*W M*_ describe how the retina-contingent stimuli move in the world. For the Gain 0 stimulus, *D*_*W M*_ and *α*_*W M*_ are equal to zero because the stimulus is not moving in the world. The world motion of the Gains *−*1.5 and +1.5 stimuli are dependent on the parameters of the eye’s motion: *D*_*EM*_ and *α*_*EM*_. How the eye moves (*α*_*EM*_) is equal to how the non-zero retina-contingent stimuli move (*α*_*W M*_) in the world. However, the eye’s magnitude of motion (*D*_*EM*_) is different from the retina-contingent stimulus’ magnitude of motion (*D*_*W M*_) because the retina contingent stimulus’s motion is transformed by a Gain. There is a squared relationship between the D and MSD, therefore, for the Gains *−*1.5 and +1.5 stimuli, the [*D*_*W M*_] = [*D*_*EM*_] *\** (1.5)^2^.

### U. Comparing the eye’s motion and the retina-contingent stimuli’s retinal motion

The *D*_*RM*_ and *α*_*RM*_ describe how the retina-contingent stimuli move across the retina. For the Gain 0 stimulus, *D*_*RM*_ and *α*_*RM*_ are equal to *D*_*EM*_ and *α*_*EM*_, respectively, because all retinal motion is the result of the stimulus slipping consistent to the fixational eye motion. The retinal motion of the Gains +1.5 and *−*1.5 stimuli are dependent on the parameters of the eye’s motion: *D*_*EM*_ and *α*_*EM*_. How the eye moves (*α*_*EM*_) is equal to how the non-zero retina-contingent stimuli move (*α*_*RM*_) on the retina. However, similar to the above paragraph the *D*_*EM*_ is not equal to the *D*_*RM*_. The Gain *−*1.5 stimulus slips with 2.5 times more motion than the eye. The Gain +1.5 stimulus moves ahead of eye motion and moves with half the motion of the eye. Therefore, for the Gain *−*1.5 stimuli, the [*D*_*RM*_] = [*D*_*EM*_] *\** (2.5)^2^; and for the Gain +1.5 stimuli, the [*D*_*RM*_] = [*D*_*EM*_] *\** (0.5)^2^.

### V. Simulated Ratios

The black line in Fig. 6 models the ratio values versus *α*_*W M*_, where the random walk stimulus’s *α* is equal to one (Brownian motion) and the retina-contingent stimulus’s *α*_*W M*_ ranged from 0.9 to 1.8. We computed the Ds where the MSDs of the random walk and retina contingent stimuli are equal. As the retina contingent stimulus’ motion becomes more persistent (*α*_*W M*_ *>* 1) the time interval over which the stimulus reaches higher MSDs becomes smaller compared to the Brownian random walk stimulus. The units of the time interval are converted to seconds because the AOSLO has a 60 Hz frame rate. The model assumes that there are no directional biases.

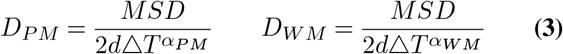

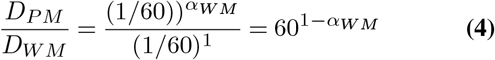

### W. Author comments

#### W.2. Author contributions

J.C.D., D.W.A, and A.R. designed research and wrote the paper; J.C.D. and A.R performed research and analyzed data; P.T. and R.W. provided critical technical support.

## ACKNOWLEDGEMENTS

We are grateful to Prof Hannah E. Smithson and Dr Allie C. Schneider for their helpful discussions. This research was supported by funding from NIH BRP R01EY023591 and NIH T32 EY007043.

